# Natural genetic variation quantitatively regulates heart rate and dimension

**DOI:** 10.1101/2023.09.01.555906

**Authors:** Jakob Gierten, Bettina Welz, Tomas Fitzgerald, Thomas Thumberger, Oliver Hummel, Adrien Leger, Philipp Weber, David Hassel, Norbert Hübner, Ewan Birney, Joachim Wittbrodt

## Abstract

The polygenic contribution to heart development and function along the health-disease continuum remains unresolved. To gain insight into the genetic basis of quantitative cardiac phenotypes, we utilize highly inbred Japanese rice fish models, *Oryzias latipes*, and *Oryzias sakaizumii*. Employing automated quantification of embryonic heart rates as core metric, we profiled phenotype variability across five inbred strains. We observed maximal phenotypic contrast between individuals of the HO5 and the HdrR strain. HO5 showed elevated heart rates associated with embryonic ventricular hypoplasia and impaired adult cardiac function. This contrast served as the basis for genome-wide mapping. In a segregation population of 1192 HO5 x HdrR F2 embryos, we mapped 59 loci (173 genes) associated with heart rate. Experimental validation of the top 12 candidate genes in loss-of-function models revealed their causal and distinct impact on heart rate, development, ventricle size, and arrhythmia. Our study uncovers new diagnostic and therapeutic targets for developmental and electrophysiological cardiac diseases and provides a novel scalable approach to investigate the intricate genetic architecture of the vertebrate heart.

**One-Sentence Summary:** Key loci for vertebrate heart function mapped and validated, highlighting diagnostic and potential therapeutic targets for cardiac disorders.

## Main Text

Cardiac phenotypes have specific morphological and functional hallmarks that can be assessed quantitatively and are typically not binary; instead, they manifest along a continuum that stretches from healthy to pathological forms. This causes a dilemma for both diagnosis and research into the early stages of the disease because normal variability between individuals examined pre-or postnatally can mask quantitative phenotypes. For example in congenital heart disease (CHD), LV hypoplasia is a quantitative phenotype ranging from mildly reduced ventricular size to a diminutive left ventricle observed in hypoplastic left heart syndrome (HLHS) (1). While the pathophysiological description of HLHS dates back to 1851 (2), its hereditary nature (3) has been a moving target as the genetics are not simple: HLHS is thought to be the outcome of multiple genetic factors that interact in an environmentally sensitive way (4, 5). Likewise, large-scale studies quantifying physiological traits with continuous individual variation, such as left ventricular parameters (6), volume measures of the right heart chambers (7), trabeculation phenotypes of the left ventricle (8), and heart rate (9, 10) indicate polygenic contribution. As a result, studies of cardiac traits have shifted from single gene analysis to exome-or genome-wide approaches. In CHD, most of these efforts have recovered private mutations (found only in one individual) and thus emphasize the polygenic nature of developmental cardiac phenotypes. Due to the limitations posed by a single genome, it has not been possible to assess how an individual’s entire collection of genomic variants drives cardiac phenotypes in processes of health and disease. Identical twins or even better isogenic lines overcome the limitations of a single genomic context and allow incorporating the variability in individual phenotypes and genotypes, environmental factors, genetic relatedness and population stratification (11–14).

Here, we turned to a well-established vertebrate model to further our understanding of genes influencing quantitative cardiac phenotypes. We use inbred strains of the teleost medaka (15) (*Oryzias latipes* (16) and *Oryzias sakaizumii* (17)) to resolve genomic variant complexity underlying quantitative cardiac trait variability. These strains represent fixed states of individually composed natural genetic variants crossed to isogenicity. They allow capturing a snapshot in the spectrum of phenotypic variation and establishing correlations with the underlying genotype by quantitative readouts in a controlled environment. To probe genetically determined cardiac phenotype variability, we used heart rate as a core phenotype, regulated by complex interactions of cardiac properties, including electrophysiology, morphology, and function. We examined the phenotypic distributions of five highly inbred medaka strains using automated heartbeat detection under controlled temperature conditions. We identified two strains, HO5 and HdrR, that differ most widely in terms of the basal heart rate. With ongoing development, the HO5 strain exhibits a hypoplastic ventricle that severely impairs fitness upon reaching adulthood. We identified 59 significant loci with a maximum association peak on medaka chromosome 3 in a quantitative trait locus mapping based on heart rate and whole-genome sequences of 1192 individually phenotyped embryos of a two-generation segregation population. We refined these loci with differential gene expression analyses in the hearts of the parental strains and teleost-to-human comparisons and validated their causal impact *in vivo* by CRISPR-Cas9 and base editor-mediated targeted gene inactivation. This demonstrated their pivotal role on heart rate, development, and morphology and revealed previously associated but functionally unrecognized loci of cardiac arrhythmia.

### Heart phenotype contrasts

The heart rate is a product of the properties of the conduction system, structural morphology, and cardiac function, and thus can be used as a readout of the way these properties are connected. We used automated microscopy and image-based heartbeat quantification under controlled physiological conditions (18), to study the maximum range of differences between phenotypes in highly isogenic inbred medaka strains/species from southern (HdrR, Cab, HO5; *O. latipes* (19)) and northern (Kaga, HNI; *O. sakaizumii* (17) Japan (Fig. 1A). We selected inbred strains/species of different geographic origins, to maximize the genetic differences and consequently cardiac phenotypes. Heart rates build up with developmental time and we addressed the dynamics of embryonic heart rates in four-hour intervals from the onset of heartbeat to the pre-hatching stage (Fig. 1B). The heart rate profiles we obtained revealed a significant and consistent spread between the five inbred strains, with the most prominent contrast between the southern *O. latipes* strains HO5 (fastest heart rate) and HdrR (slowest heart rate, Fig. 1B).

**Fig. 1.**
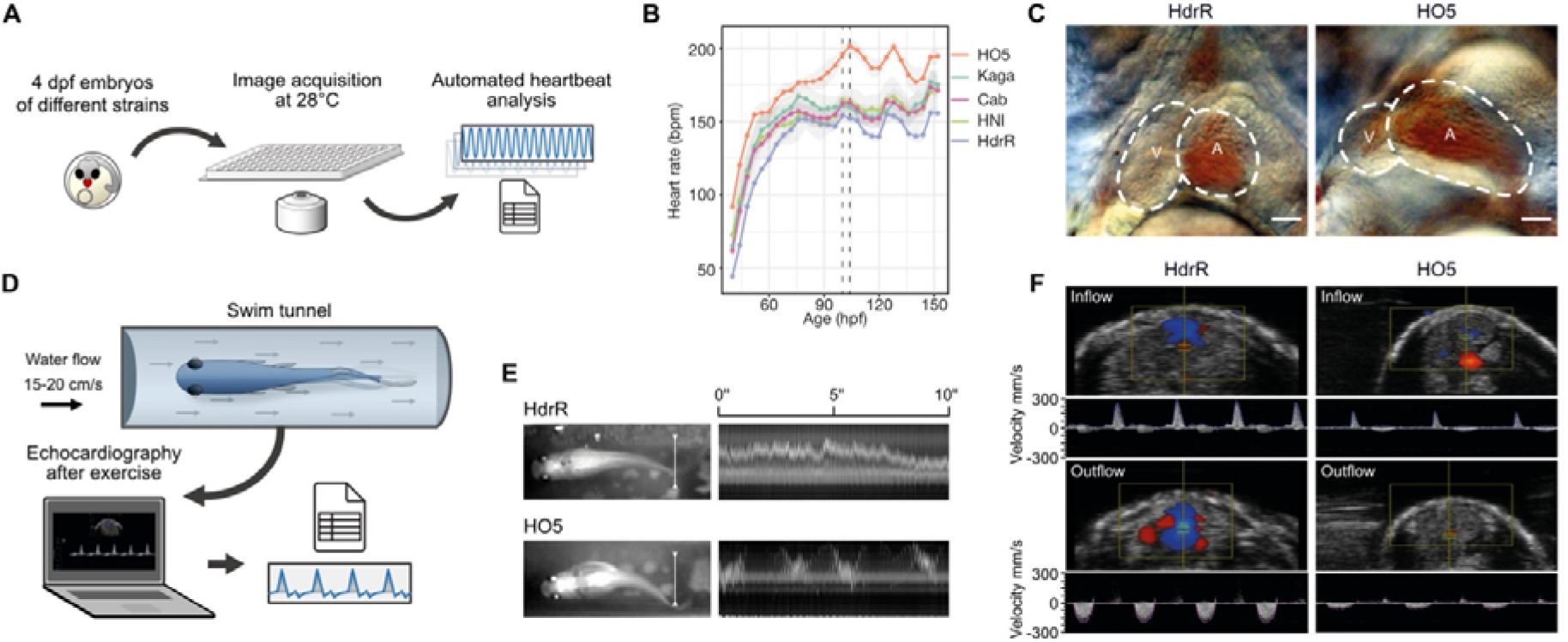
Cardiac phenotype contrast in medaka inbred strains. (**A**) Layout of the automated heartbeat detection in medaka embryos in native environment (28°C) using high-throughput imaging and image-based heart rate quantification. (**B**) Distribution of embryonic heart rates in five inbred strains derived from Southern Japanese medaka populations (HdrR, HO5, Cab) and Northern Japanese populations (HNI, Kaga) across embryonic development starting with the onset of heartbeat. Heart rates of 6-18 embryos per strain were determined every four hours under a 12 h-light/12 h-dark cycle; smoothed line between single (4 h interval) heart rate measurements for each strain (table S1); dotted lines, window of circadian-rhythm-stable heartbeat for comparative analysis (100-104 hours post fertilization, hpf). (**C**) Cardiac morphology in HdrR and HO5 hatchlings; end-systolic frame, scale bar, 50 µm. (**D, E**) Exercise assessment and swim performance of adult fish in a swim tunnel assay (movie S1). White line in (E) used for kymograph - note stable swimming behavior in HdrR versus fluctuating HO5 individual. (**F**) Pulsed-wave (pw) doppler of ventricular inflow (atrium-ventricle) and outflow (ventricle-bulbus arteriosus) tracts.

We ensured consistency by measuring heart rates in a narrow time window between 100-104 hpf (dotted rectangle, Fig. 1B) after the completion of critical stages of cardiac development (20) and thus avoided the impact of circadian oscillations observed from 3 dpf onwards (18).

The differences in embryonic heart rate were substantial, prompting us to examine cardiac morphology at 4 dpf (raised at 28 °C). HO5 embryos exhibited unbalanced heart chambers with a dominant atrium and an underdeveloped (hypoplastic) ventricle. This apparent hypoplastic ventricular morphology was associated with a high basal heart rate (Fig. 1C), potentially compensating a reduced ventricular output. As the unique combination of natural genetic variation fixed in the HO5 genome contributes to its cardiac phenotype, we next assessed the extent to which it influences the function and physical fitness of the heart at adult stages. We subjected adult HO5 and HdrR individuals to a swim tunnel exercise protocol (Fig. 1E). Video monitoring indicated the efficient performance of HdrR; fish assumed stable positions in a defined water flow, requiring minimal fin excursions to generate the necessary swimming speed (Fig. 1F and movie S1). In contrast, HO5 individuals (apparently higher BMI) used almost the entire body length to generate forward movement. In addition, they failed to assume stable positions within the constant water flow (cf. kymograph in Fig. 1E). Following swim tunnel exercise, we found that HO5 exhibited significantly reduced velocities in intracardiac blood flow, indicating reduced myocardial function, as observed using echocardiography, including pulse-waved (pw) Doppler measurements of the ventricle in- and outflow (Fig. 1F). These findings argue for early alterations of physiological cardiac traits that occur in HO5 embryos and severely impact on physical fitness and cardiovascular health in adulthood.

### Segregation analysis

To map the loci contributing to these complex phenotypes, we established a mapping population in which we correlated heart rates and whole-genome sequences (WGS). To model a human-relevant modifiable environmental factor of embryonic development, we turned to the well-established heart rate increase in elevated water temperatures(18) and measured embryonic heart rates at an incremental ambient temperature ramp (21°C, 28°C, 35°C). For mapping we employed a two-generation segregation design and used single-nucleotide polymorphisms (SNPs) as markers (Fig. 2A). SNP calling in HO5 WGS against the HdrR reference genome established 979,713 differential homozygous SNPs as marker system.

**Fig. 2.**
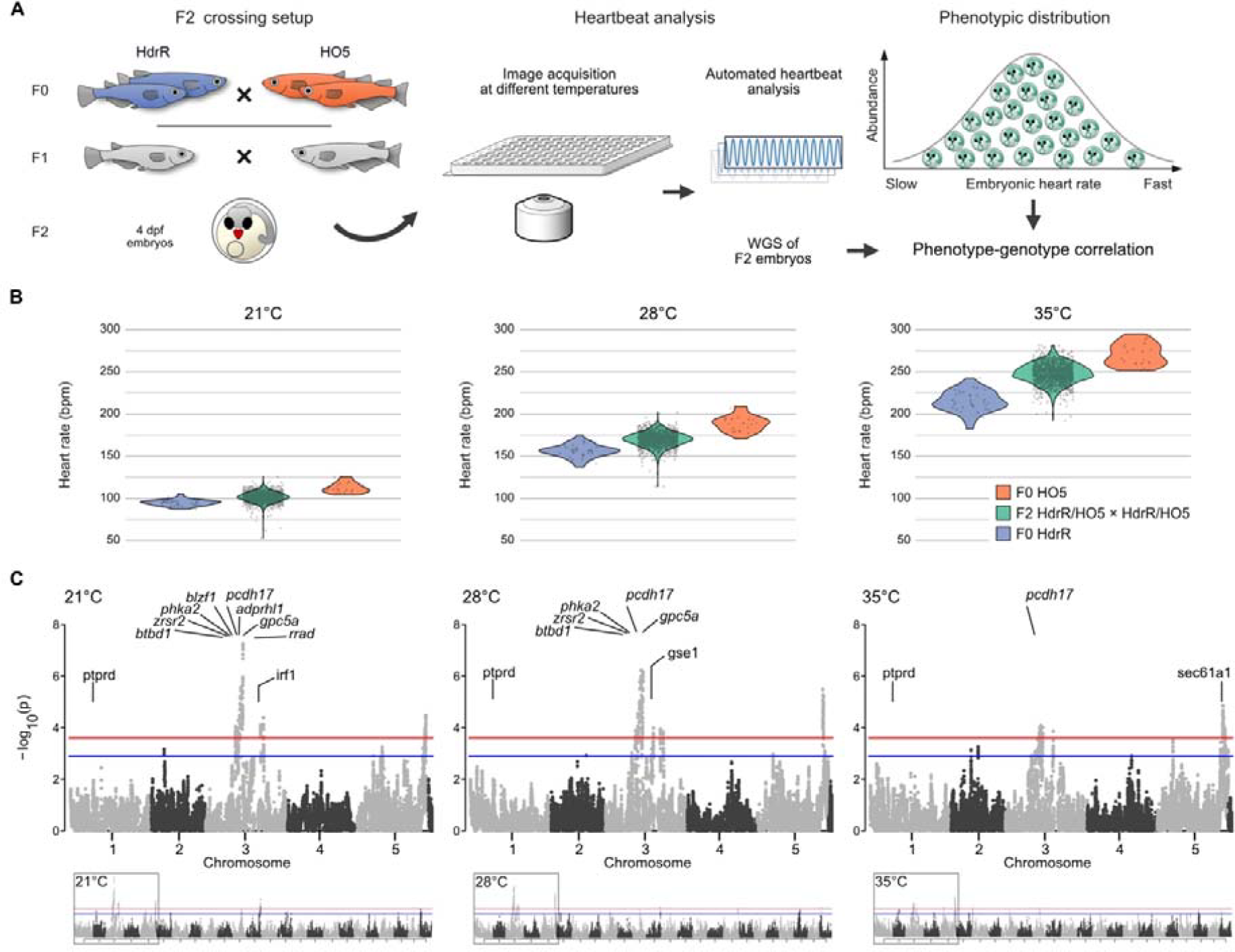
F2 segregation analysis reveals temperature-sensitive QTLs affecting the heartbeat. (**A**) Crossing setup used to generate HdrR × HO5 offspring with segregated SNPs in the second generation: isogenic HO5 and HdrR parents are crossed to generate hybrid (heterozygous) F1 generation (grey) with intermediate phenotype, which after incrossing results in F2 individuals with individually segregated SNPs resulting from one cycle of meiotic recombination. Automated embryonic heartbeat analysis at 4 days post fertilization (dpf), performed on all 1260 individual embryos and challenged by increasing temperatures (21 °C, 28 °C, 35 °C). Whole-genome sequencing of measured individuals (1192 genomes) with an effective average coverage of 0.78× allows phenotype-genotype correlation. (**B**) Individual embryonic heart rates of the inbred strains HdrR (slow heart rate, blue) and HO5 (fast heart rate, orange) increase with temperature (21 °C, 28 °C and 35 °C). The F2 individuals with recombined HdrR × HO5 genomes (green) span the range of parental (F0) heart rates between the two strains with a subgroup of F2 individuals exhibiting heart rate variance beyond the parental extremes (21 °C and 28 °C); sample sizes (n) for 21 °C, 28 °C and 35 °C: n (HdrR F0) = 35, 35, 35, n (HO5 F0) = 20, 22, 22, n (HdrR × HO5 F2) = 1260, 1260, 1260. (**C**) Minus log10 p values from genome-wide association tests of recombination block genotypes and heart rate measures at different temperatures using a linear mixed model. Chromosomes 3 and 5 hold the most segregated recombination blocks associated with heart rate differences. Twelve selected genes from the loci passing the significance threshold are indicated; red line: 1% false discovery rate (FDR), blue line: 5% FDR, determined by permutation.

We crossed the two isogenic strains with distinctly differentiated cardiac phenotypes to generate a hybrid F1 population with haploid sets of chromosomes from each parental strain and observed heart rates that were intermediate between the two parental strains. We next performed F1 intercrosses and sampled 1260 individuals of the F2 generation with unique inherited genotypes (linkage blocks) to serve as the mapping population (Fig. 2A). We conducted individual heart rate phenotyping of all 1260 F2 embryos and found (with a few outliers showing sporadic arrhythmia) that the phenotypic distribution covered the range between the parental phenotypes at all three temperatures (Fig. 2B). This reflects the differential segregation of multiple alleles that directly impact the heart rate or secondarily due to morphological-functional effects on the heart.

### Genome-wide QTL mapping

We raised individually 1260 phenotyped F2 embryos in 96-well plates followed by automated DNA extraction, and whole-genome-sequencing. We sequenced 1192 samples with an average coverage of 0.78×. We used a three-state Hidden Markov Model (HMM) to segment all crossover locations and determine genotype states (AA, AB, BB) based on SNPs homozygous divergent in HO5 (AA) and HdrR (BB). Interestingly, we observed a distortion in the expected Mendelian ratio of 1:2:1 (AA:AB:BB alleles) with an overrepresentation of AA (HO5) alleles suggestive of a potential reference bias. This bias was evident in most F2 offspring samples (fig. S1) and was not restricted to specific regions of the genome. Although it could be possible that certain crossover events between HdrR and HO5 are incompatible, the most parsimonious explanation is a tendency towards homozygous reference calls within the SNP genotype calls used to train the HMM. Having called crossover events and generated a recombination map across all F2 offspring samples independently, we merged the crossover locations and segmented the genome at every breakpoint resulting in a genotype matrix containing AA:AB:BB calls for variable sized blocks. The median genotype block size, once segmented across all F2 cross recombination positions, was 24 kb.

Using this genome matrix we then conducted genome-wide association analyses of 101,265 segmented recombination block regions on individual heart rate measures at three different ambient temperatures and for absolute differences in repeated measurements across all temperatures 21°C, 28°C and 35°C (variance phenotype), using a linear mixed model from the GridLMM software package (21). We found significantly associated quantitative trait loci (QTLs) that were mostly consistent across the three temperatures (Fig. 2C) and different peaks in the test on heart rate variance from 21-35°C (fig. S2). Overall, we detected 1385 significant loci across all phenotype tests and performed collapsing down to 59 distinct fine-mapped regions after linkage disequilibrium (LD)-based SNP pruning. The maximum achievable resolution for loci fine mapping was 10kb due to the window size used for recombination block mapping, and the median size across all fine-mapped loci was 115 kb. We were able to fine-map to 10 kb for only 5/59 loci; however, most fine-mapped regions contained small numbers of genes (median of 1 and a range between 0-36 genes per fine-mapped block). Overall, we detected similar numbers of fine-mapped loci for the different phenotype measures we tested, with 17, 16, 8, and 17 unique fine-mapped regions for 21°C, 28°C, 35°C and the variance-based phenotype, respectively.

To aid gene prioritization and interpretation of significantly associated loci in the F2 population, we undertook total RNA sequencing of heart tissues in 4 samples from both the HO5 and HdrR strains. After quality control 18,321 genes (75% of known medaka genes) had sufficient coverage to allow differential expression analysis. At a false discovery rate (FDR) of 0.01 and minimum fold change of 2 we detected 1,161 significantly differentially expressed genes between HO5 and HdrR hearts (fig. S3A), which we then used to prioritize genes within significantly associating F2 recombination blocks for subsequent validation experiments. Genes expressed in liver samples of both strains were used as base line reference (fig. S3B). Additionally we assessed the likely impact of SNP calls in HO5 against the HdrR reference using the variant effect predictor (VEP) from Ensembl (22). As expected, loss-of-function (LoF) variants were rare within these blocks with a median of 0 variants and maximum of 3 per gene. However using the combination of information, it was possible to rank genes and variants more likely to be impactful for the observed phenotypic differences. One case supported by LoF variation is a single premature stop codon variant in the *blzf1* gene, and in contrast the *rrad* gene where no LoF variants in HO5 were observed but a significant differential expression in heart tissues.

### Gene enrichments

Across all 59 significantly associated fine mapped regions there were 173 annotated genes in the *Oryzias latipes* (HdrR) ENSEMBL reference, 28 of the fine mapped regions had no annotated gene within their boundaries (8, 10, 6 and 4 loci with no annotated genes for heart rate phenotypes at 21°C, 28°C, 35°C, and variance respectively), leaving 34 containing 1 or more genes with a median of 3 per fine mapped region. The largest region was found on chromosome 1 associated with the variance phenotype and contained 36 genes, only two of which had any LoF variant called in HO5 against the HdrR reference (data S1, block ID-2). Using the 173 annotated genes, we performed multiple gene enrichment models using the PANTHER overrepresentation test (released 20230705), gProfiler functional profiling and functional annotation clustering using DAVID (23–25). Using the overrepresentation test from PANTHER, we found significant associations (FDR P < 0.05) for 72 different biological process GO terms including for “thrombocyte differentiation”, which has a greater than 100-fold enrichment. Both gProfiler and DAVID found a significant enrichment (adjusted P value 4.393×10-10) for the GO term “galactoside binding” with 6 of the 172 genes being involved in the binding of glycoside carbohydrate derivatives, as well as a weaker but significant enrichment (adjusted P value 1.895×10-2) for the KEGG term “Linoleic acid metabolism”. For functional annotation clustering using DAVID with a classification stringency set to Medium using the Benjamini adjustment, there were 21 clusters defined, the strongest of which had an overall enrichment score of 4.56 and included the GO terms “galactoside binding” and “laminin binding”.

Recent studies in humans have looked at the pleiotropic regulatory activities of Galectins in relation to cardiovascular disease (CVD) and their proinflammatory role in the atherosclerotic and plaque formation process (26). Here we found 6 Galectin related genes (ENSORLG00000030479, ENSORLG00000026315, ENSORLG00000010700, ENSORLG00000010715, ENSORLG00000010697, ENSORLG00000024256) to be associated to heart rate differences in medaka, associated to the variance in heart rate and found within the same fine mapped region at chromosome 8:13655000-13935000 (Data S2, block ID-42). Galectins are promiscuous, with multiple cellular functions and impede arteriogenesis (27). Consequently, galectin inhibitors are an attractive target for therapeutics pertaining to remodeling in myocardial infarction (28). Thus, associations based on medaka heartbeat dynamics directly surfaced genes with highly relevant translational implications.

### *In vivo* validation

Our genome-wide QTL mapping based on heart rate metrics identified loci associated with biological functions in heart development, cardiac function, and electrophysiology. To provide evidence for a causal role of the identified loci, we selectively deactivated genes in medaka embryos and investigated the respective impact on embryonic heart rate and morphology *in vivo*. After analyzing the search space (table S2), we narrowed our selection to 12 candidate genes based on their high linkage probabilities, differential expression, human relevance, and novelty (cf. Methods “Candidate gene selection”): *adprhl1, blzf1, btbd1, gpc5a, gse1, irf1, pcdh17, phka2, ptprd, rrad, sec61a1*, and *zrsr2*, where *rrad* and *adprhl1* have previously been directly associated with cardiovascular traits.

We addressed potential functional effects on the heart following targeted inactivation of candidate genes by CRISPR-Cas9-or base editor-mediated knockout. Phenotype analysis was performed after cardiovascular development was expected to be complete at 4 dpf (28°C) in embryos devoid of global developmental defects, which resulted in two major phenotypes: embryos with morphologically normally developed hearts and cardiac-specific affected embryos, showing looping defects, pericardial edema, arrhythmia, or aberrant atrial or ventricular size and morphology (Fig. 3, fig. S4, table S3). Cas9-based targeted inactivation of candidate genes prominently increased the proportion of cardiac phenotypes above the baseline threshold established by injected controls (max. 7% heart phenotypes; Fig. 3B and fig. S4). Specifically, *adprhl1, blzf1, btbd1, phka2, ptprd*, and *rrad* showed high rates of prominent heart phenotypes in the CRISPR-Cas9 approach. To underline the specificity of observed phenotypes, we corroborated the results of the CRISPR-Cas9 gene interrogations in an independent experimental series introducing premature termination codons (PTCs) with the cytosine base editor evoBE4max for six of the twelve selected candidate genes (Fig. 3C).

**Fig. 3.**
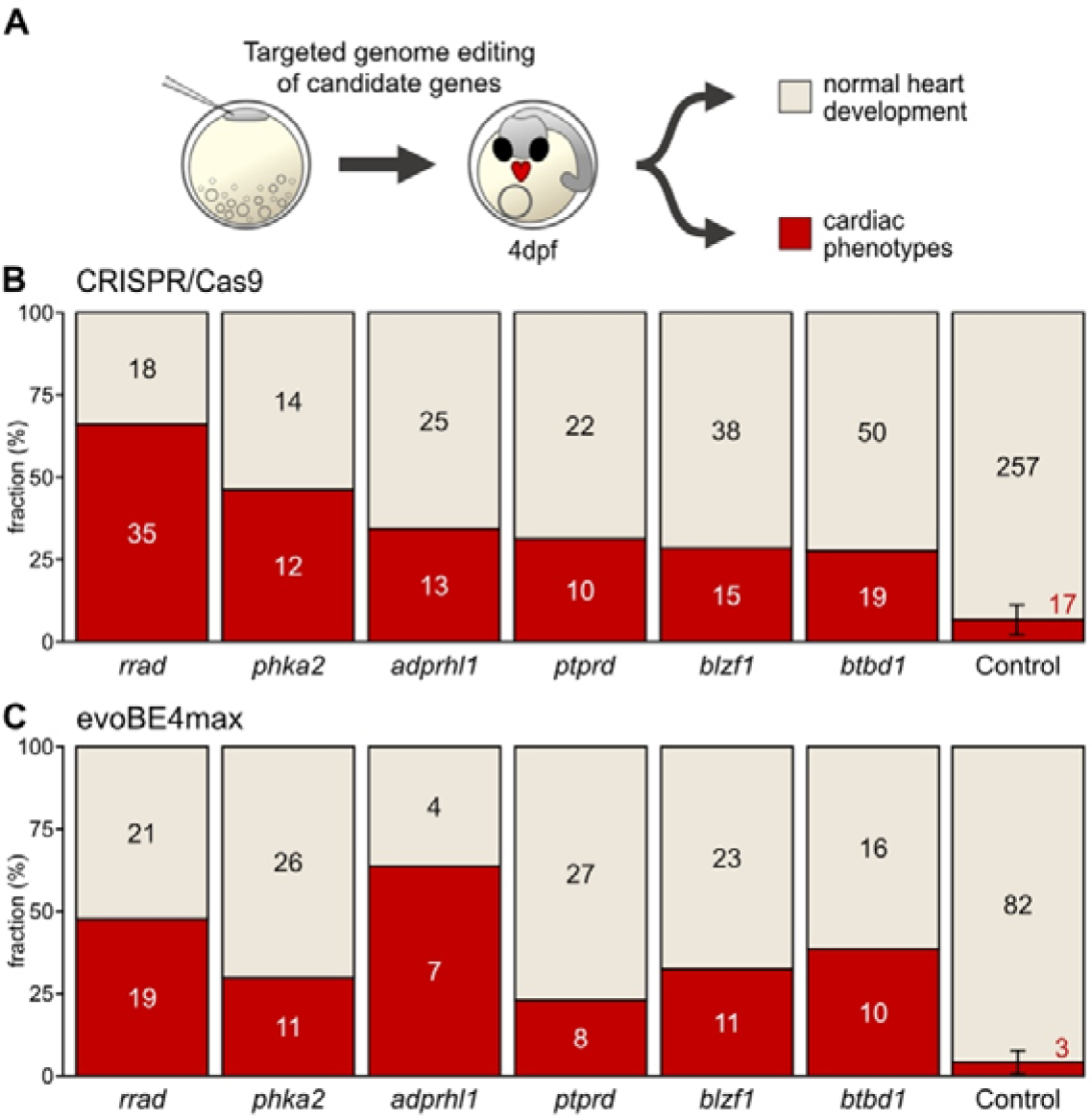
Phenotype proportions in knockout models of cardiac candidate genes. (**A**) Validation workflow encompassing zygotic microinjections using a HdrR (*myl7::eGFP*; *myl7::H2A-mCherry*) reporter line, followed by phenotypic classification of embryos with normal developed hearts and embryos with cardiac-specific phenotypes 4 days post fertilization (dpf). (**B**) Proportion (bars) and counts (values) of cardiac affected and normally developed embryos after CRISPR-Cas9-mediated knockout of indicated candidate genes versus control (mock injection). (**C**) Independent replication using base editing: phenotypic distribution (proportion/bars and counts/values) resulting from targeted gene inactivation mediated by introducing premature termination codons via the cytosine base editor evoBE4max and a set of distinct guide RNAs targeting the same genes as in (B).

To discriminate potential specific effects of candidate genes on heart rate and exclude secondary effects in embryos with aberrant cardiac morphogenesis, we applied the automated heart rate assay in embryos with apparently normal heart development (Fig. 4, fig. S5). We found that for eight out of the twelve candidate genes, the heart rate was affected (data S2). While six of them showed an increase in heart rate across the three different temperatures tested in replicates with Cas9- and base editor evoBE4max-mediated targeted gene inactivation (Fig. 4B and C), interfering with *zrsr2* and *gse1*, conversely, resulted in a marked decrease in heart rate (fig. S5).

**Fig. 4.**
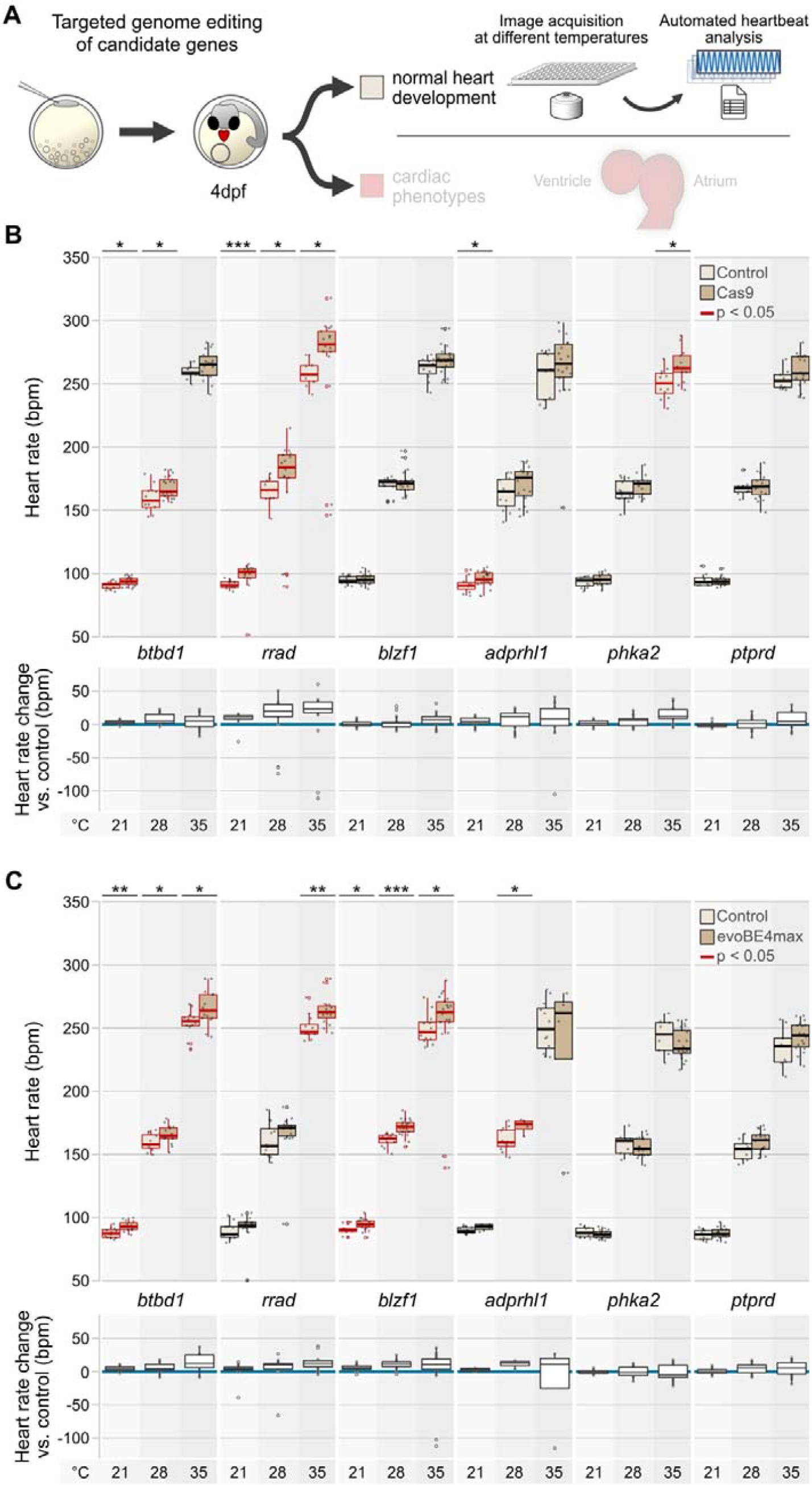
Loss-of-function mutations in candidate genes affect embryonic heart rate in medaka embryos. (**A**) Validation workflow including zygotic microinjections using a HdrR (*myl7::eGFP*; *myl7::H2A-mCherry*) reporter line, high-throughput image-based heart rate quantification in normally developed injected specimens at 4 days post fertilization (dpf). (**B, C**) Heart rate distributions and absolute heart rate differences of morphologically normal mutants compared to mock-injected control embryos at 21°C, 28°C, and 35°C with CRISPR-Cas9-(B) and base editor-mediated (C) targeted mutagenesis of candidate genes. The significance of heart rate differences between the mutant group and its corresponding control was assessed with the Wilcoxon test; *, p<0.05; **, p<0.01; ***, p<0.001 (p-values listed in table S3). Data is visualized as box plots (median+/-interquartile range) and overlaid scatter plots of heart rate measurements; knockouts of *adprhl1, btbd1, blzf1, phka2*, and *rrad* have temperature-dependent effects on heart rate.

Our automated heartbeat quantification algorithm is optimized to extract heartbeats from cardiac segments with the most robust image signal, not discriminating between the atrial and ventricular frequencies. Consequently, this heartbeat quantification uncovers arrhythmias as apparent as marked outliers in the case of *rrad, adprhl1*, and *blzf1* (Fig. 4B and C). Manual examination of the corresponding image series revealed embryos with aberrant atrioventricular conduction and block, resulting in a marked drop in the automatically scored ventricle beat rate.

We further closely assessed the morphological aberrations and functional cardiac specific phenotypes (quantified in Fig. 3) and the gene inactivations triggering arrhythmia (observed in Fig. 4B and C) (Fig. 5, Fig. S6). While targeted inactivation of *ptprd, blzf1*, and *btbd1* distorted cardiac looping (incorrect positioning of the atrium to the right and the ventricle to the left), a reduced ventricle size was observed in specimen after targeted inactivation of *rrad, adprhl1, blzf1*, or *btbd1* (Fig. 5A). Heartbeat detection surfaced cardiac conduction defects in crispants of three candidate genes (*rrad, blzf1, adprhl1*; Fig. 4B and C). Interestingly, these genes have been previously associated with electrophysiological phenotypes and disorders. While *blzf1* had been associated with cardiac repolarization (QT interval) (29), *rrad* with the specific human cardiac conduction disorder Brugada syndrome (30), and *adprhl1* with left anterior fascicular block (31), these associations had not been experimentally validated. We re-assessed the phenotype in compound heterozygous F2 embryos and verified the initially observed AV block with 2:1 conduction in the case of *rrad* and *adprhl1*, while *blzf1* mutants displayed an irregular lack of AV conduction (Fig. 5C).

**Fig. 5.**
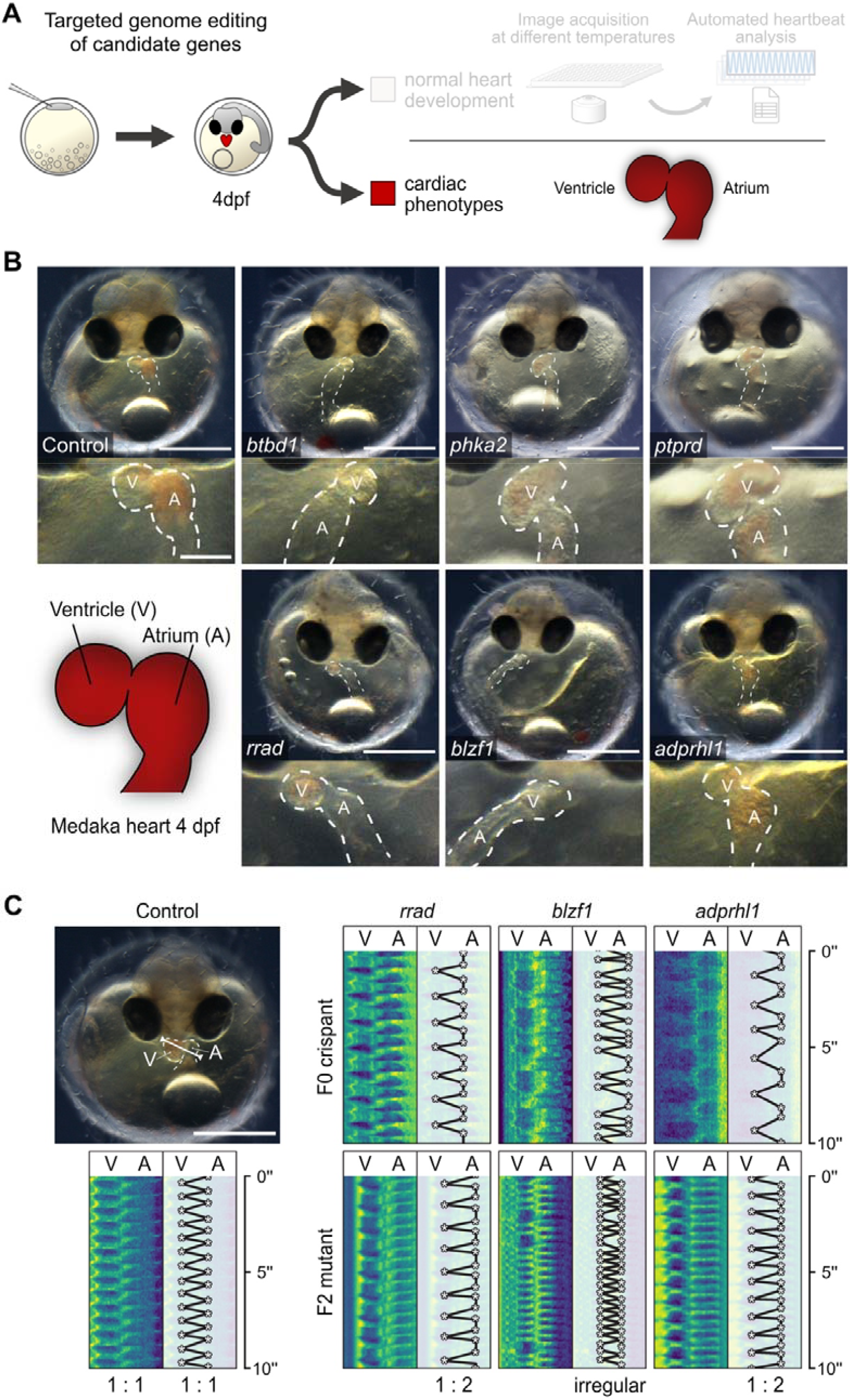
Developmental and electrophysiological phenotypes in F0 crispants and F2 mutants. (**A**) Workflow including zygotic microinjections using a HdrR (*myl7::eGFP*; *myl7::H2A-mCherry*) reporter line and phenotyping of cardiac affected crispants 4 days post fertilization (dpf). (**B**) Representative cardiac phenotypes of knockout embryos for all six candidate genes targeted with CRISPR-Cas9 and base editor compared to a control embryo at 4 dpf; bright-field overview of the injected specimen (top; scale bar, 500 µm), close-up image of the heart (bottom; scale bar, 125 µm); cf. movie S2. (**C**) Heart rhythm analysis in F0 crispants and F2 mutants as depicted by kymographs derived from atrium (A) to ventricle (V) spanning line selection in 10 second time-lapse movies. In contrast to the regular rhythm in the control embryo (left), the representative *rrad, blzf1 and adprhl1* F0 crispant and F2 mutants (right) display different degrees of atrioventricular block; cf. movie S4.

Taken together, the F2 gene segregation workflow combined with quantitative phenotyping and targeted mutagenesis for validation allowed to robustly uncover novel genetic risk factors for human heart disease.

## Discussion

Using highly inbred medaka strains, we mapped specific SNPs that contribute to differences in heart rate, rhythm, and ventricular size disproportion, phenotypes interrelated in human CHD. Analyzing almost 1200 genomes and the corresponding heart rate as readout with predictive power (9, 10), we identified 59 QTLs at high resolution. 173 of the genes contained in these mapped loci showed a direct link to heart as well as extracardiac organ functions, reflecting the polygenic architecture of quantitative heart phenotypes.

We specifically inactivated a subset of these genes to validate their potential role in the development of cardiac phenotypes, based on a differential expression analysis in medaka and an orthology-guided examination of genes associated with human heart phenotypes. We carried out CRISPR-Cas9 and base-editing-mediated inactivation of these candidates and uncovered genes involved in heart rate and rhythm control as well as structural development of the heart. Intriguingly, using heart rate metrics for phenotype-genotype mapping, we identified genes with dual roles affecting heart muscle mass and rhythm; e.g., *rrad* combining regulative functions on cardiac muscle strength and electrophysiology. *Rrad* is known to inhibit cardiac hypertrophy through the CaMKII pathway with implications for heart failure (32), and is associated with hypertrophic cardiomyopathy (HCM) phenotype in RRAD-deficient cell line (33). *Rrad* mutations detected in a specific form of familiar arrhythmia (Brugada syndrome) trigger cytoskeleton and electrophysiological abnormalities in iPSC-CMs (30), which we validated *in vivo*, demonstrating a specific atrioventricular block in medaka *rrad* crispants, base-edited embryos and mutants. Additionally, we found the heart rate-associated genes *blzf1*, located on a locus associated with cardiac repolarization but previously not recognized as a functional candidate (29), and *adprhl1*, associated with a ventricular conduction disorder (31), producing specific conduction defects in our generated knockout models. This demonstrates the power of complex cardiac trait analysis in medaka inbred strains to detect human-relevant association signals, guiding genome editing experiments to uncover novel players and potential targets in heart disease.

We noticed a significant decrease in ventricular size in a subset of gene knockouts (*rrad, adprhl1, blzf1*, and *btbd1*), and it appears that *adprhl1* has a major impact on ventricular dimensions. Adprhl1 is a conserved pseudoenzyme with ADP-ribosylarginine hydrolase and magnesium ion binding activity, likely exclusive to the heart. Transcriptomics revealed very strong cardiac expression (3,109.244 RPM compared to weak (1.175 RPM) in the liver) in medaka. Until now, *adprhl1* has been studied functionally only scarcely. Of note, in *Xenopus, adprhl1* can localize to the cardiac sarcomeres. While morpholino-mediated knockdown of *adprhl1* did not affect early cardiogenesis, it disrupts myofibril assembly in the forming ventricle and leads to small, inert ventricles (34). In light of the hypoplastic ventricle observed in the HO5 strain, its location on a strongly associated QTL on chromosome 3 in medaka, and targeted inactivation *in vivo*, we provide evidence for *adprhl1* as a strong contender for controlling myofibril assembly and ventricular outgrowth.

The homozygous fixation of causative genomic variants in viable inbred medaka strains allows not only for modeling twin studies with arbitrary scalability but also for longitudinal investigation to estimate the variants’ effects within a lifespan. Notably, we found an inverse relation of metrics for embryonic heart rate and ventricle size in the HO5 strain, which exposes pathological heart phenotype with impaired cardiac function and physical performance in exercise tests, demonstrating the predictive power of genotype-specific embryonic heart rate profiles. Finding such early biosignatures is essential for diagnosing and potentially preventing severe pathophysiologies that become less likely correctable over the long term.

Most cardiac disease phenotypes are quantitative and manifest in a health-disease continuum. Here, we demonstrate that inbred medaka represents a powerful resource to partition and validate the spectrum of phenotypes in a strain-specific way. Examining the extremes of a physiological trait range bears the power to reveal the genetic factors likely associated with disease susceptibility. The F2 cross of inbred strains with contrasting phenotypes allows to segregate causal alleles in a way that cannot be achieved with human GWAS studies. Nevertheless, the results can be immediately transferred to the human context and shed light on genetic variations that have flown under the radar.

In summary, our research underscores the invaluable role of inbred medaka strains in unraveling the intricacies of cardiac development, physiology, and pathology. The comprehensive genetic analysis conducted here has not only led to the discovery of novel heart-related genes but also established a promising, robust framework for future investigations.

By leveraging the power of a scalable medaka inbred panel (35–37) in conjunction with highly reliable quantitative phenotyping assays across the entire phenotypic spectrum, as further detailed in the accompanying manuscript by Seleit et al., we anticipate a crucial enhancement in our ability to pinpoint and validate combinatorial candidate variants across both coding and non-coding regions of the genome. This holds a great perspective for advancing our understanding of cardiac biology in particular and may pave the way for innovative therapeutic interventions in cardiovascular disease.

## Supporting information

biorxiv combined supplements

Supplementary Table 2 - mapping results

Supplementary Table 3 - heart beats

## Acknowledgments

We thank Steffen Lemke and Lazaro Centanin for critically reviewing the manuscript and the Wittbrodt, Birney, Lemke and Centanin labs for constructive support of the work and manuscript. Thanks to R. Hodge for critically reading and commenting on the manuscript. We thank J. Gehrig (ACQUIFER, Bruker) for technical support in high-throughput imaging and benchmarking algorithms. We also thank T. Kellner, B. Wittbrodt, and R. Müller for excellent technical support as well as, E. Leist, M. Majewski, A. Saraceno and S. Erny for fish husbandry.

## Funding

European Research Council (ERC) Synergy grant 810172/IndiGene (EB, JW)

National Institutes of Health (NIH) grant R01ES029917 (EB, JW)

German Centre for Cardiovascular Research (DZHK) grant 81X2500189 (JG, JW, NH)

Deutsche Herzstiftung e.V. grant S/02/17 (JG)

Research Center for Molecular Medicine (HRCMM), Career Development Fellowship (JG)

Medical Faculty Heidelberg University, MD/PhD program (JG)

Joachim Herz Stiftung, Add-On Fellowship for Interdisciplinary Science (JG)

## Author contributions

Conceptualization: JG, EB, JW

Investigation: JG, BW, TF, PW, TF, EB, AL, EB, OH, NH, JW, TT

Visualization: JG, BW, TF, TT, AL

Funding acquisition: JG, EB, JW, NH

Project administration: EB, JW, DH

Supervision: DH, EB, JW, NH

Writing - original draft: JG, BW, TT, JW, TF, EB

