## Supplementary material for "Natural genetic variation quantitatively regulates heart rate and dimension": biorxiv combined supplements

Joachim Wittbrodt^1,3^*

†These authors contributed equally to this work

**The PDF file includes:**

Materials and Methods

Figs. S1 to S6

Tables S1 to S7

References (38-50)

**Other Supplementary Materials for this manuscript include the following:**

Movies S1 to S4

Data S1 to S2

**Materials and Methods**

Fish maintenance

The wild-type Kaga, HNI, Cab, HO5 and HdrR strain and a fluorescent cardiac reporter line HdrR (*myl7::EGFP myl7::H2A-mCherry*) were used in this study. All fish stocks were maintained (fish husbandry, permit number 35–9185.64/BH Wittbrodt) and experiments (permit numbers 35– 9185.81/G-145/15, 35–9185.81/G-10/17 and 35–9185.81/G-271/20) were performed in accordance with local animal welfare standards (Tierschutzgesetz §11, Abs. 1, Nr. 1) and with European Union animal welfare guidelines (38). Fish were kept as described previously (39). The fish facility is under the supervision of the local representative of the animal welfare agency.

Automated microscopy and heartbeat detection of medaka embryos

The automated microscopy and heart rate quantification was applied as described previously (18). Embryonic heart rate profiles over developmental stages were generated by imaging embryos in 96-well plates every four hours under a 12 h-light-12 h-dark-cycle, i.e., 12 h dark minus imaging time of 20 min/plate/time point; incubation temperature was 28 °C (developmental profiles) and 21°C, 28°C, and 35°C in the screen. For display “smoothed conditional means'' encoded in ‘geom_smooth()’ using regression method = ‘loess’ (ggplot2) with 95% confidence interval was used.

All F0, F1 and F2 embryos were imaged as separate batches in 96-well plates each with one row (12 embryos) of Cab embryos as an internal control. For phenotype-genotype correlations in individual F2 embryos, fifteen 96-well plates, each containing two F2 crosses (HdrRf × HO5m F2 and HO5f × HdrRm F2), were imaged and processed under the same conditions.

To exclude positional effects of plate coordinates on heart rate, a full 96-well plate with Cab embryos was recorded at 28°C. Heart rates were normally distributed (*P*=0.86; Shapiro-Wilk normality test). One-way ANOVA indicated no significant effects of plate row on mean heart rates (degrees of freedom (df) = 7.88; *P* = 0.61), nor of plate column on mean heart rates (df = 11.84; *P* = 0.15).

Swim tunnel assay and echocardiography of adult medaka fish

Two months before the experiments, HdrR F96 and HO5 F112 were kept at 22°C water temperature and 14 h/10 h light/dark conditions. Swim tunnel (Loligo Systems) assay was performed with the following flow velocity and settings. Equilibration at 3 cm/s for 20 min, then increase of water flow by 5 cm/s every 5 min; mO2 was scored at each step. Two 10 s videos with 100 fps were recorded at 15 cm/s and 20 cm/s (if applicable) during the interval after a flush. The test was stopped when fish remained 3 s or more at the rear of the chamber. Echocardiography was performed as previously described (40) with 150 mg/l tricaine and 8 mg/l metomidate.

HdrR × HO5 intercross and phenotyping of F0, F1, and F2 embryos

For heart rate measurements in F0 (i.e., HO5 F112, HdrR F95, Cab F68), embryos were collected from these crosses: 3 female × 1 male HO5 F111, 3 female × 1 male HdrR F94 and as plate control 3 female × 1 male Cab F67.

Phenotyping F1 hybrid embryos and generation of F1 hybrid stocks: 3 HO5 F111 females × 1 HdrR F94 male resulting in stock HO5 F111f × HdrR-II F94m F1, and 3 HdrR F94 × 1 HO5 F111m resulting in stock HdrR F94f × HO5 F111m F1. Cab F68 embryos were used as plate control obtained from 3 female × 1 male Cab F67.

The embryonic F2 screen (heart rate) was conducted within two months. Two F2 populations were derived from two separate crosses of each F1 hybrid stock (1) HO5 F111f × HdrR F94m F1 and

(2) HdrR F94f × HO5 F111m F1: F2 embryos were collected from stocks (1) and (2), for each of which 3 tanks with 3 females × 1 male were set up (both F2 populations derived each from 12 fish). Cab F68 embryos were used as plate control obtained from 3 female × 1 male Cab F67. Incubation conditions for F0, F1 embryos: medaka embryos were raised at 28°C with either max. 20 embryos/20 ml medaka hatching medium in 60 mm dishes or with max. 50 embryos/50 ml medaka hatching medium in 90 mm dishes.

Incubation conditions for F2 embryos: 55 embryos were cultured at 28°C in 90 mm dishes with medaka hatching medium; Cab controls: 25 embryos in 9 cm dishes with hatching medium. Embryo preparation for imaging: Embryos were rolled on sandpaper the afternoon/evening before imaging and placed back to the primary culture dishes with fresh hatching medium.

Heart rate assay (cf. “Automated microscopy and heartbeat detection of medaka embryos”): On the day of imaging, embryos were transferred from hatching medium to ERM before mounting. Individual F2 embryos were loaded with 150 µl ERM in 96-well plates (U-bottom). Approximate imaging times were: Start equilibration at 12:00, imaging 21°C at 12:15 PM, 28°C at 13:15 PM and 35°C at 14:15 PM.

F2 sample preparation and whole genome-sequencing

After imaging of F2 embryos in 96-well plates, ERM was exchanged with medaka hatching medium by pipetting out 140 µl ERM and adding 200 µl medaka hatching medium. Subsequently, 190 µl of the medaka hatching medium was exchanged daily until first embryos hatched. When the majority of embryos had hatched, the remaining unhatched embryos were manually dechorionated. All hatchlings were transferred from the 96-well imaging plate to identical coordinates of a 96-DeepWell plate, which was covered with an adhesive aluminum foil (4titude) and frozen acutely at -80°C until further processing. Samples were processed at Wellcome Sanger Institute (UK).

Library preparation and WGS on a HiSeq X Ten instrument (Illumina) with an average sequencing depth of 0.78×. Library preparation was performed following the standard PCR-free Illumina protocol (41). In total, 1 μg was picked from DNA extraction plates within the Sanger sample logistics facility and passed into the Sanger sequencing pipeline from library preparation. Following successful preparation of sequencing libraries, the samples were QCed using Qubit and samples passing the facility quality control threshold were multiplexed sequenced in paired end mode with 150 samples per Illumina X10 flow cell.

Whole genome-sequence analysis and mapping

For the genotyping of recombination blocks in the F2 population, a modified algorithm of the previous version was used (https://github.com/tf2/ViteRbi). The analysis was based on SNPs homozygous divergent in HO5 (AA) and HdrR (BB). Homozygous divergent SNPs were determined by the alignment of high-depth HO5 genome sequencing data against the HdrR-II reference genome. At each homozygous divergent SNP-locus read counts supporting each genotype (A, T, C, G) were extracted from the genome alignments. In a fixed window of 5000 bp, read counts were summed and transformed into proportions of reads supporting the AA genotype. This genotype proportion was the observational input for a three-state Hidden Markov Model employing the Viterbi algorithm to segment all crossover locations and to determine genotype states (AA, AB, BB) in the resulting recombination blocks across all samples. Genome-wide association tests using recombination block genotypes and heart rate measures were performed using a linear mixed model (21).

RNA sequencing and transcriptome analysis

Heart and liver samples were dissected from euthanized adult HO5 F112 and HdrR F96 female fish and collected into Qiagen Collection Microtubes (racked, 19560) with cabs (19566), acutely frozen on dry ice and then stored at -80°C. Total RNA purification was performed with QIAsymphony RNA, purifying liver samples with RNA CT 400 and heart samples with RNA FT 400. Samples were prepared for Illumina RNA sequencing using the NEBNext Ultra II Directional RNA Library Prep Kit for Illumina and sequenced on a Hiseq 4000 sequencing platform following the manufacturer’s instructions. All pair-end reads were cleaned up using Fastp v0.20.0 (42), aligned to the medaka HdrR reference genome Ensembl Release 98 (ASM223467v1) using STAR v2.7.3a (43) and estimated counts per gene were obtained. Differential expression analysis was performed using DESEQ2 (44) with an FDR cut-off of 0.01 and fold change of 2, 1161 and 1803 significantly differentially expressed genes were obtained for heart and liver samples respectively.

Candidate gene selection

QTLs were prioritized according to their strength of linkage. To filter the search space defined in Data S2 we included recombination blocks (QTLs) with SNP-based genotypes significantly (log10 p-value>3) associated with heart rate variation. Only the strongest linkage to heart rate variation is given for each block compared at three temperatures (21°, 28°, and 35°C). Phenotype associations of orthologous human genes supported candidate gene selection relevant to human cardiac disease. Therefore, the human GWAS genome-wide association studies (GWAS) catalog (gwas_catalog_v1.0.2-associations_e104_r2021-10-22.tsv downloaded from https://www.ebi.ac.uk/gwas/docs/file-downloads) (45) was scanned for terms indicative of heart rate and morphology associations. Candidate genes were additionally studied with the GeneALaCart batch query processor of the GeneCards suite (https://www.genecards.org) (46). If blocks contain multiple genes, candidates were chosen according to defined criteria, primarily novelty and human relevance, positively (+) or negatively (-) ranking a candidate gene (table S1).

sgRNA and crRNAs target site selection

rrad (ENSORLG00000024517) sgRNAs, adprhl1 (ENSORLG00000004693) sgRNAs, ptprd (ENSORLG00000004685) sgRNAs, phka2 (ENSORLG00000003555) sgRNAs, blzf1

(ENSORLG00000003670)	sgRNAs,	btbd1	(ENSORLG00000003434)	sgRNAs,	zrsr2

(ENSORLG00000003476) sgRNA, pcdh17 (ENSORLG00000004535) sgRNA, sec61a1 (ENSORLG00000016830) sgRNA, irf1a (ENSORLG00000011716) sgRNA, gse1 (ENSORLG00000009542) sgRNA, and gpc5a (ENSORLG00000026952) sgRNA were designed with CCTop and ACEofBASEs as described previously (39, 47), on the medaka genome in Ensembl release 101 (Japanese medaka HdrR assembly, Aug 2020), Ensembl release 102 (Japanese medaka HdrR assembly, Nov 2020), Ensembl release 103 (Japanese medaka HdrR assembly, Feb 2021) and Ensembl release 106 (Japanese medaka HdrR assembly, Apr 2022), respectively. The target sites and oligonucleotides selected for sgRNA cloning are listed in tables S5 and S6. Cloning of sgRNA templates and in vitro transcription was performed as described previously(47). The plasmid DR274 used was a gift from Keith Joung (Addgene 42250). For base editing, locus-specific crRNAs and the tracrRNA backbone were obtained from IDT (custom AltR crRNA). crRNA (100 µM) and tracrRNA (100 µM) were diluted in nuclease-free duplex buffer (IDT) to a final concentration of 40 µM and incubated at 95°C for 5 min.

In vitro transcription of mRNA

The plasmid pCS2+(Cas9) and pCS2+(evoBE4max) were linearized using NotI, and mRNA in vitro transcription was performed using the mMESSAGE mMACHINE SP6 or T7 Transcription Kit (Thermo Fisher Scientific) and purified with the RNeasy Mini Kit (Qiagen).

Microinjections

Microinjections were performed in HdrR (*myl7::EGFP*, *myl7::H2A-mCherry*) zygotes. The Cas9 injection solution contained 150 ng/µl Cas9 mRNA,10 ng/µl sgRNA and 10 ng/µl GFP mRNA as injection tracer. The base editor injection solution contained 150 ng/µl evoBE4max mRNA, 4 pmol sgRNA and 10 ng/µl GFP mRNA as injection tracer. Control injections were performed with 10 ng/µl GFP mRNA as injection tracer. Injected embryos were maintained at 28°C in medaka embryo rearing medium (ERM, 17 mM NaCl, 40 mM KCl, 0.27 mM CaCl2, 0.66 mM MgSO4, 17 mM Hepes) and selected for GFP expression 7 hours post injection. Phenotypes were assessed 4 days post fertilization.

Image acquisition and phenotyping of knockout models

Embryo morphology and heart dynamics were documented with a Nikon SMZ18 equipped with Nikon DS-Ri1 and DS-Fi2 cameras. Embryos were mounted in EMR into injection molds (1.5 % (w/v) agarose in ERM). Heart rate quantification of CRISPR-Cas9 and evoBE4max-mediated knockout embryos and control embryos were obtained using an ACQUIFER Imaging Machine and the HeartBeat software as described previously (18, 48). To acquire the heart rate, fluorescent image sequences have been captured for 10 sec with 24 fps at 21°C, 28°C, and 35°C.

Genotyping

Single phenotypic, non-phenotypic and control embryos were lysed in DNA extraction buffer (0.4 M Tris/HCl pH 8.0, 0.15 M NaCl, 0.1% SDS, 5 mM EDTA pH 8.0; 1 mg/ml proteinase K) at 60°C overnight. Samples were diluted 1:2 with nuclease-free water and the proteinase K was heat inactivated afterwards for 20min. Precipitation of genomic DNA was done in 300 mM sodium acetate and 3x vol. absolute ethanol at 20,000 x g at 4°C, followed by resuspension in TE buffer (10 mM Tris pH 8.0, 1 mM EDTA in RNAse-free water). The target loci were PCR amplified in 30 PCR cycles using Q5 High-Fidelity DNA Polymerase (New England Biolabs), locus-specific primer pairs (table S7) and 1 µl precipitated DNA sample. PCR products were purified after agarose gel electrophoresis using the Monarch DNA Gel Extraction Kit (New England Biolabs) and submitted for Sanger sequencing to Eurofins Genomics. Base editing results were analyzed with EditR (49).

Data analysis and statistics

Data visualization and statistical analysis were performed using R (50)

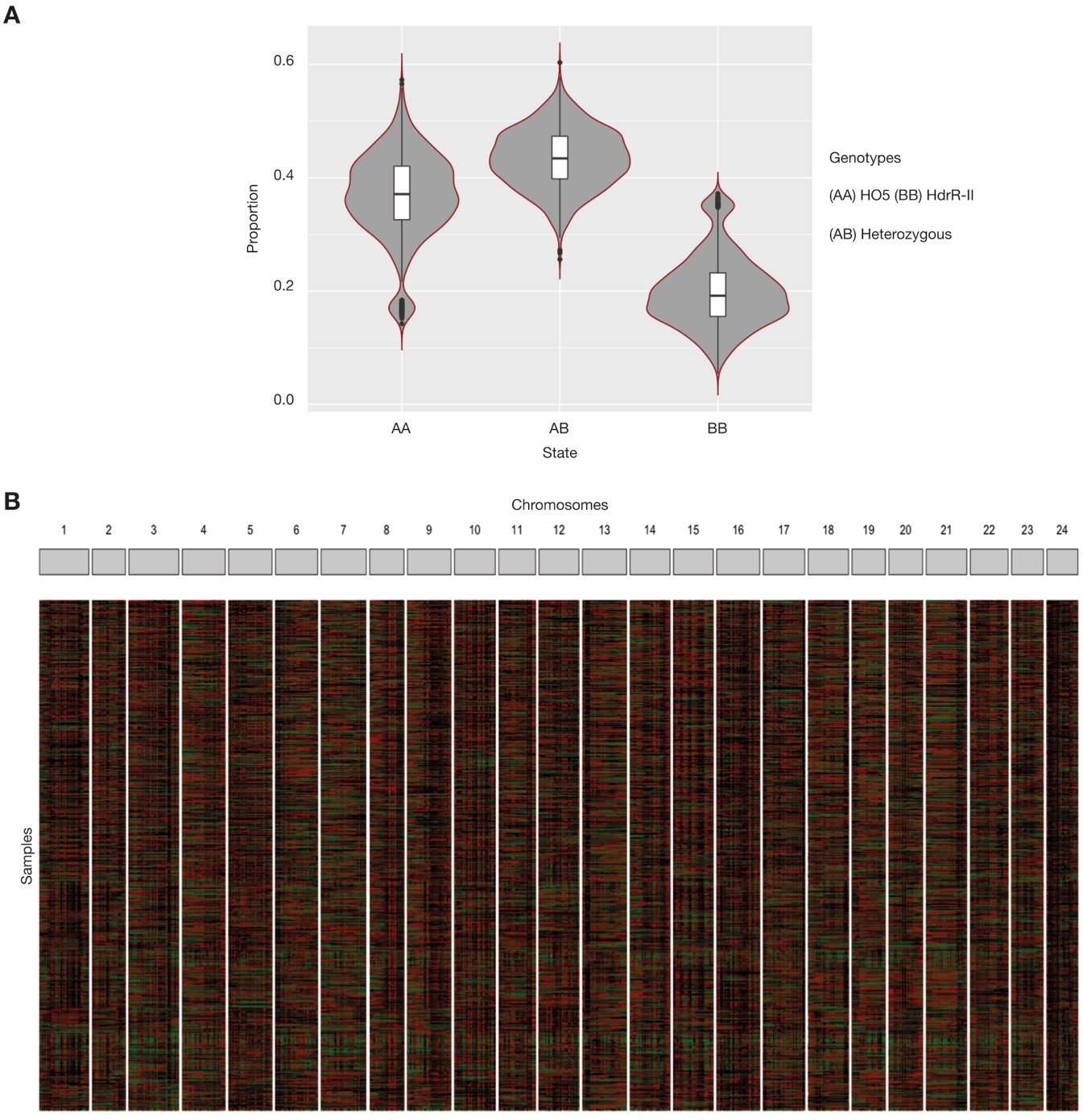

**Fig. S1 Genotype proportions of the parental lines HdrR and HO5 in the sequenced F2 population.** (**A**) Proportions of homozygous HO5 genotype (AA), homozygous HdrR genotype (BB) or heterozygous genotype (AB). (**B**) Genotypes and recombination block distribution for the F2 samples (rows) across the 24 chromosomes (columns). AA = black, BB = green, AB = red.

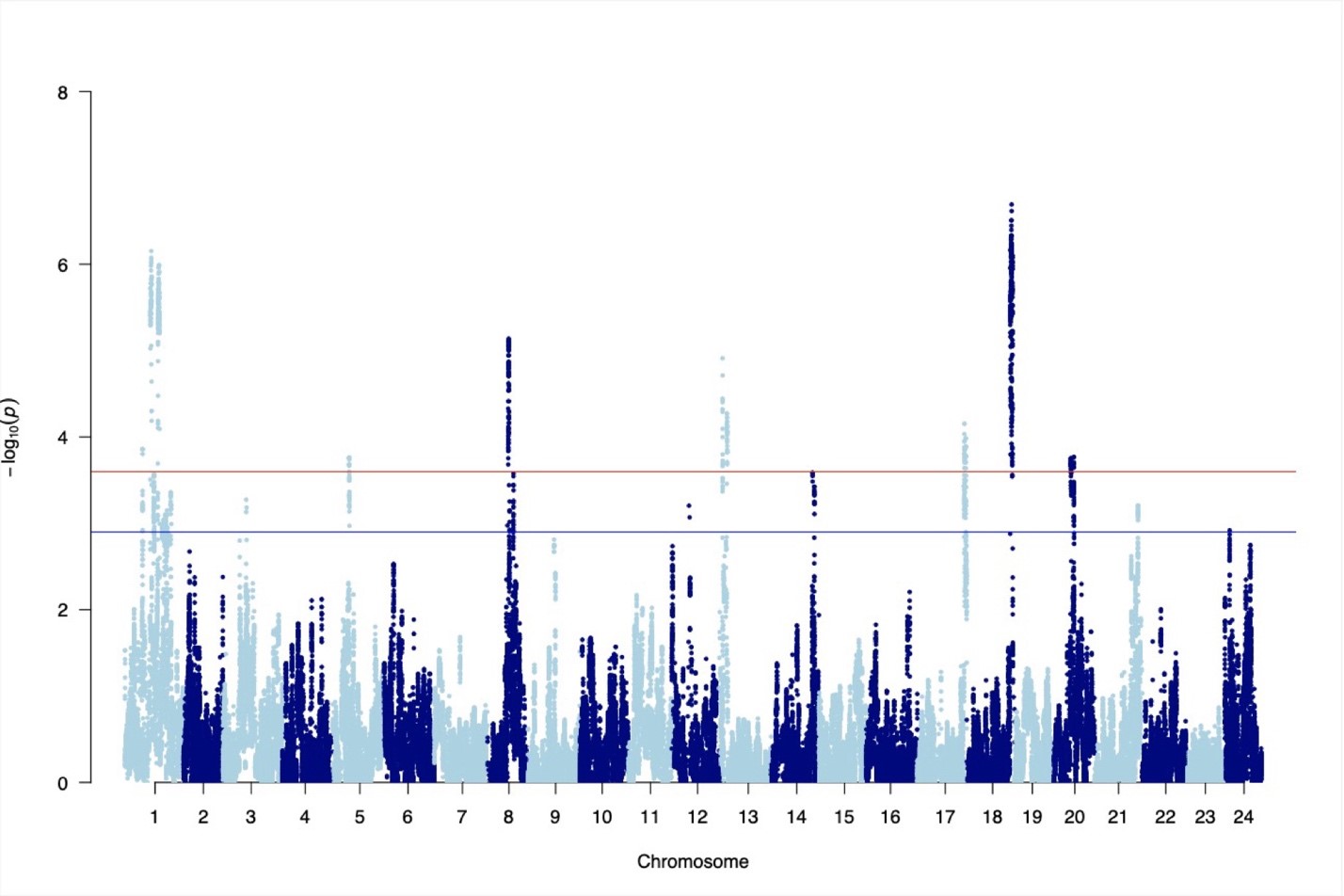

**Fig. S2 Variance phenotype.** Manhattan plot showing -log10 p values from the linear mixed model using the variance phenotype (mean absolute difference between repeated measurements on the same embryo across the 3 different temperatures).

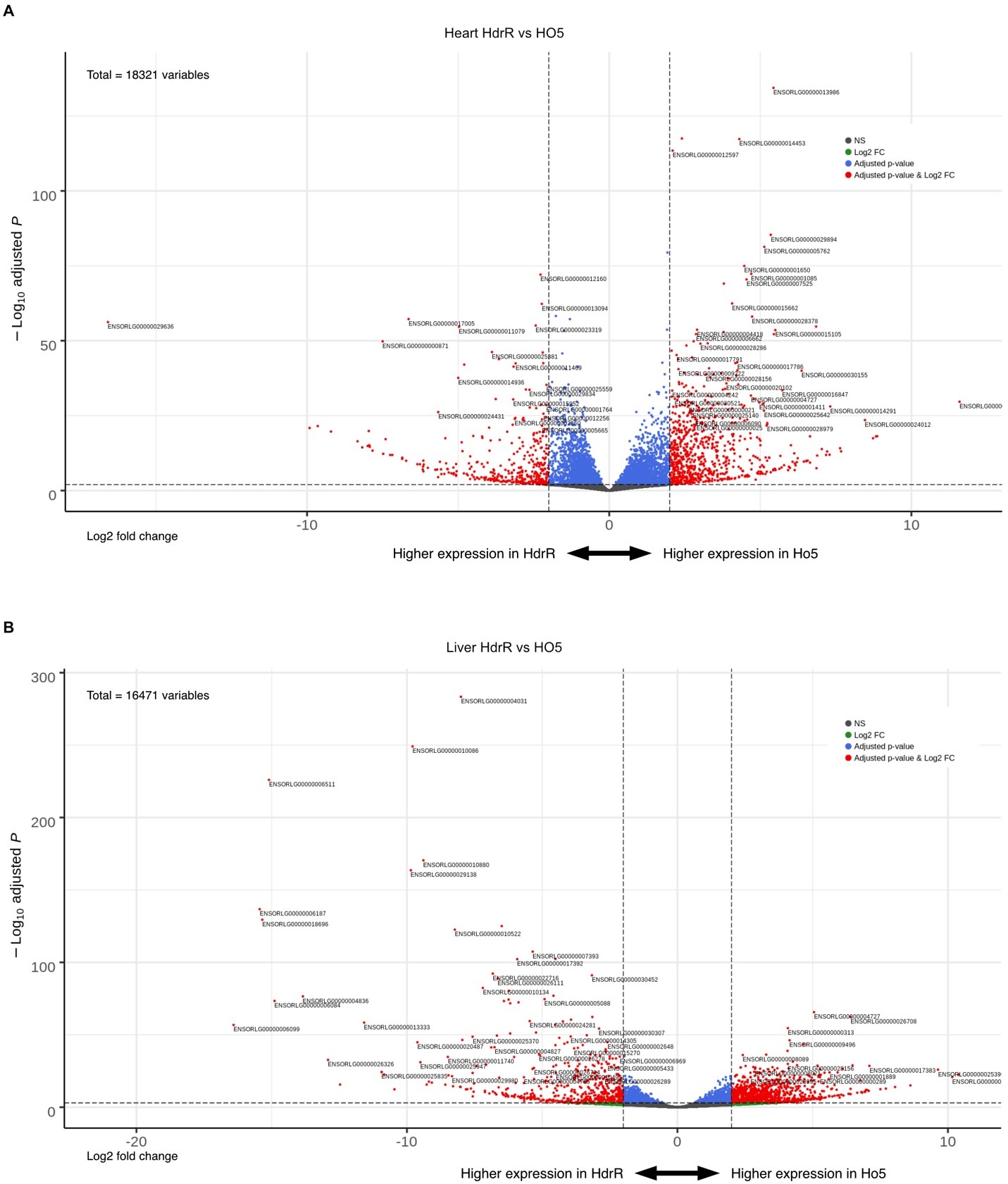

**Fig. S3 Differentially expressed genes in HdrR versus HO5 heart and liver samples.** (**A**, **B**) Volcano plots of HdrR and HO5 heart (A) and liver (B) comparative transcriptomics, -log10(adjpvalue) compared with log2 Fold change.

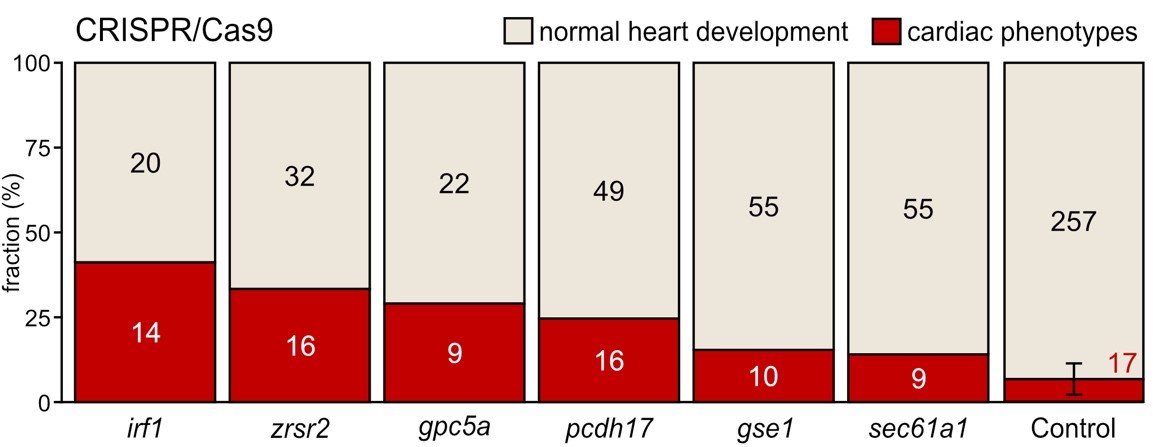

**Fig. S4 Heart phenotype proportions in CRISPR-Cas9 knockout models of candidate genes.** (**A**) Proportion (bars) and counts (values) of cardiac affected and normally developed embryos after CRISPR-Cas9-mediated knockout of indicated candidate genes versus control (mock injection) quantified at 4 days post fertilization (dpf).

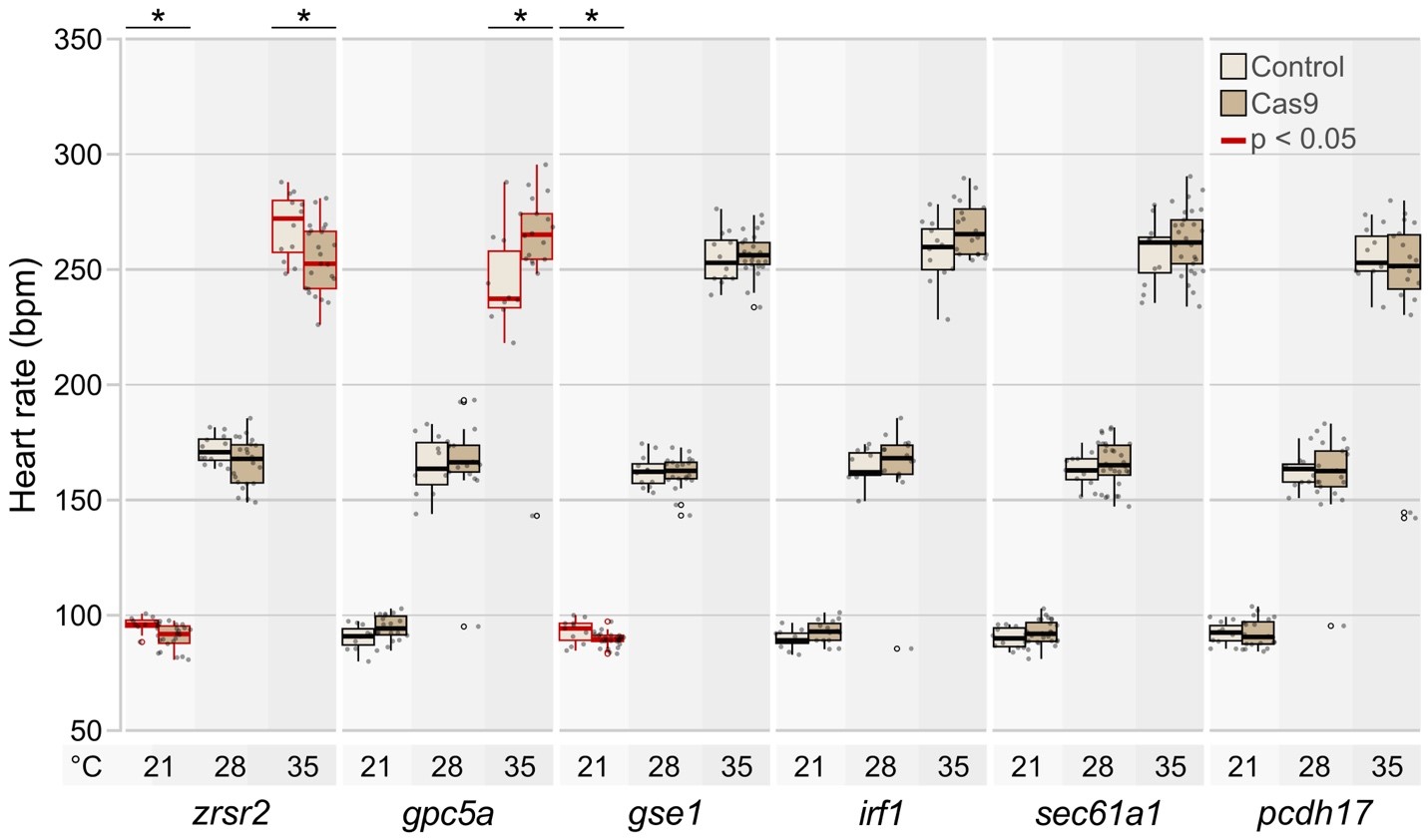

**Fig. S5 Loss-of-function candidate gene mutations affecting the heart rate level of medaka embryos.** Heart rate distributions of morphologically normal CRISPR-Cas9-mediated gene lossof-function embryos and mock-injected control embryos at 4 dpf at 21°C, 28°C, and 35°C. The significance of heart rate differences between the mutant group and its corresponding control was tested with the Wilcoxon test; *, p<0.05, **, p<0.01, ***, p<0.001 (p-values for all comparison groups are listed in Table S4). Data is visualized as box plots (median+/- interquartile range) and overlaid scatter plots of heart rate measurements; *zrsr2* mutants and *gse1* mutants show a temperature-dependent significant decrease in heart rate, in contrast, *gpc5a* mutants show a significant increase in heart rate at 35°C.

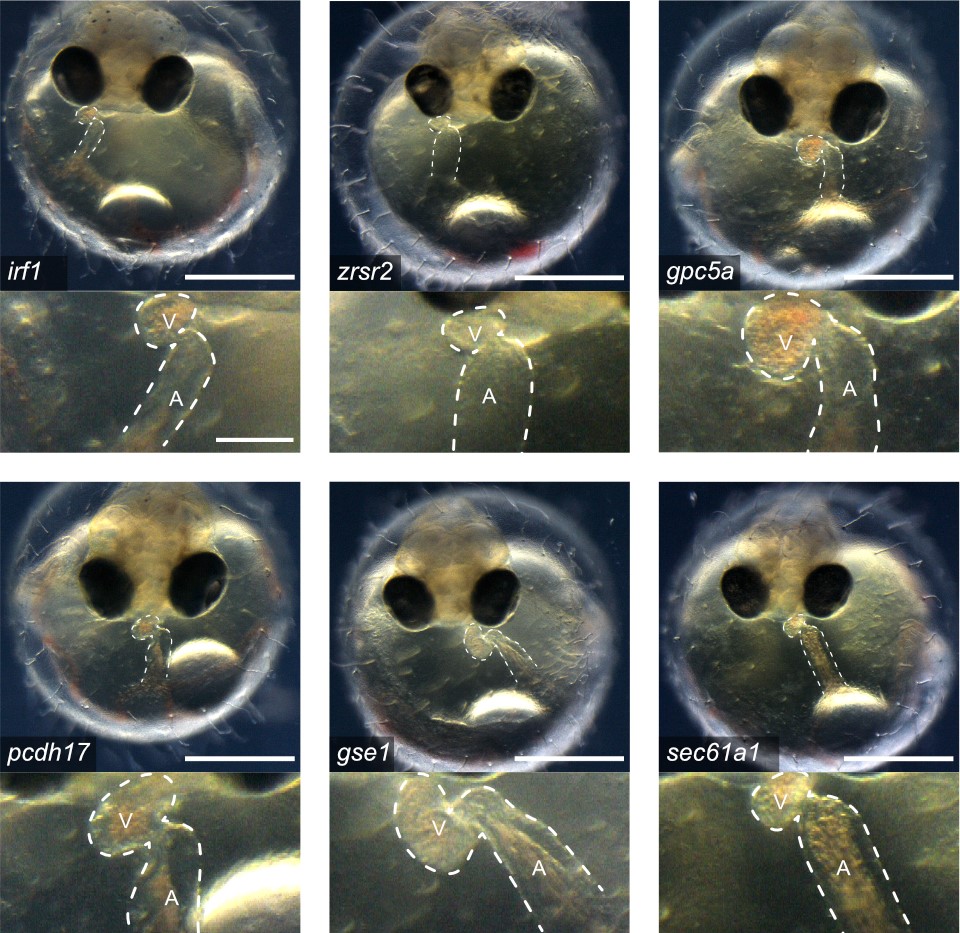

**Fig. S6 Morphological heart phenotypes in CRISPR-Cas9 knockout models of candidate genes.** Cardiac phenotypes of knockout embryos for the six candidate genes targeted with CRISPR-Cas9 at 4 dpf; bright-field overview of the injected specimen (top; scale bar, 500 µm), close-up image of the heart (bottom; scale bar, 125 µm). cf. movie S3.

**Table S1 Numbers of embryos scored (n) in Fig. 1B for each strain and point in time. Fluctuations in the sample sizes are due to random embryonic movements during imaging that can obscure the heart and preclude measurement; embryos that have already hatched (swimming) have been excluded resulting in a drop of n at 148 hpf and 152 hpf.**

| **Age (hpf)** | **n HdrR** | **n HNI F86** | **n HO5** | **n Cab** | **n Kaga** |
| --- | --- | --- | --- | --- | --- |
| 40 | 14 | 14 | 18 | 15 | 15 |
| 44 | 18 | 17 | 18 | 18 | 15 |
| 48 | 18 | 17 | 18 | 17 | 15 |
| 52 | 18 | 18 | 18 | 18 | 15 |
| 56 | 18 | 18 | 18 | 18 | 15 |
| 60 | 18 | 18 | 17 | 18 | 15 |
| 64 | 18 | 17 | 18 | 18 | 15 |
| 68 | 18 | 18 | 18 | 18 | 15 |
| 72 | 18 | 18 | 18 | 18 | 15 |
| 76 | 18 | 18 | 16 | 18 | 15 |
| 80 | 18 | 16 | 12 | 16 | 14 |
| 84 | 18 | 18 | 18 | 18 | 15 |
| 88 | 18 | 18 | 16 | 17 | 15 |
| 92 | 18 | 18 | 15 | 17 | 13 |
| 96 | 18 | 18 | 18 | 18 | 15 |
| 100 | 18 | 17 | 15 | 18 | 15 |
| 104 | 18 | 13 | 16 | 16 | 10 |
| 108 | 16 | 16 | 17 | 17 | 15 |
| 112 | 18 | 15 | 18 | 18 | 15 |
| 116 | 17 | 14 | 17 | 18 | 15 |
| 120 | 18 | 15 | 16 | 17 | 15 |
| 124 | 17 | 14 | 17 | 17 | 11 |
| 128 | 14 | 13 | 18 | 16 | 14 |
| 132 | 17 | 13 | 18 | 17 | 11 |
| 136 | 18 | 13 | 18 | 18 | 14 |
| 140 | 16 | 12 | 16 | 18 | 13 |
| 144 | 18 | 10 | 18 | 18 | 13 |
| 148 | 16 | 10 | 18 | 18 | 7 |
| 152 | 15 | 9 | 17 | 18 | 6 |

**Table S2 Workflow of candidate gene prioritization.**

| **Ranking criterion** | **Database** | **Positive (+) or negative () influence on gene rank** |
| --- | --- | --- |
| (1) Differential expression in the heart | Transcriptomes, this study | + |
| (2) Presence of a paralog in medaka  (Oryzias latipes) | Ensembl | - |
| (3) Known cardiac phenotype in medaka  (Oryzias latipes) | Ensembl | - |
| (4) Presence of a high confidence  ortholog in zebrafish (Danio rerio) | Ensembl | + |
| (5) Known cardiac phenotype in zebrafish | Ensembl, ZFIN | - |
| (6) Known cardiac phenotype in mouse or rat | Ensembl, MGI/JAX | - |
| (7) Presence of high confidence ortholog in human | Ensembl | + |
| (8) Cardiovascular > ubiquitous  expression | Ensembl, NCBI | + |
| (9) (a) Presence of ortholog or (b) variant(s) associations in human | Ensembl | + |

**Table S3 Number (n) of injected and phenotyped embryos at 4 dpf. Surviving embryos were grouped into three categories: normal, cardiac affected and global phenotypes.**

| **Gene** | **sgRNA/crRNA** | **n injected** | **n normal** | **n**    **cardiac** | **n global** | **n dead** | **Editor** |
| --- | --- | --- | --- | --- | --- | --- | --- |
| rrad | rrad_T1 sgRNA | 93 | 18 | 35 | 20 | 20 | CRISPR_Cas9 |
| adprhl1 | adprhl1_T1 sgRNA | 81 | 25 | 13 | 17 | 26 | CRISPR_Cas9 |
| btbd1 | btbd1_T1 sgRNA | 100 | 50 | 19 | 21 | 10 | CRISPR_Cas9 |
| blzf1 | blzf1_T1 sgRNA | 84 | 38 | 15 | 14 | 17 | CRISPR_Cas9 |
| ptprd | ptprd_T1 sgRNA | 54 | 22 | 10 | 8 | 14 | CRISPR_Cas9 |
| phka2 | phka2_T1 sgRNA | 62 | 14 | 12 | 7 | 29 | CRISPR_Cas9 |
| pcdh17 | pcdh17_T1 sgRNA | 92 | 49 | 16 | 12 | 15 | CRISPR_Cas9 |
| irf1 | irf1a_T1 sgRNA | 83 | 20 | 14 | 27 | 22 | CRISPR_Cas9 |
| gpc5a | gpc5a_T1 sgRNA | 72 | 22 | 9 | 25 | 16 | CRISPR_Cas9 |
| gse1 | gse1_T1 sgRNA | 79 | 55 | 10 | 4 | 10 | CRISPR_Cas9 |
| sec61a1 | sec61a1_T1 sgRNA | 81 | 55 | 9 | 8 | 9 | CRISPR_Cas9 |
| zrsr2 | zrsr2_T1 sgRNA | 114 | 32 | 16 | 43 | 23 | CRISPR_Cas9 |
| rrad | rrad_T2 crRNA | 75 | 21 | 19 | 15 | 20 | evoBE4max |
| adprhl1 | adprhl1_T2 crRNA | 68 | 4 | 7 | 14 | 43 | evoBE4max |
| btbd1 | btbd1_T2 crRNA | 53 | 16 | 10 | 10 | 17 | evoBE4max |
| blzf1 | blzf1_T2 crRNA | 68 | 23 | 11 | 13 | 21 | evoBE4max |
| ptprd | ptprd_T2 crRNA | 54 | 27 | 8 | 0 | 19 | evoBE4max |
| phka2 | phka2_T2_crRNA | 50 | 26 | 11 | 0 | 13 | evoBE4max |
| mock_ctrl1 | NA | 41 | 36 | 1 | 0 | 4 | NA_Cas9_ctrl |
| mock_ctrl2 | NA | 49 | 44 | 3 | 0 | 2 | NA_Cas9_ctrl |
| mock_ctrl3 | NA | 46 | 42 | 0 | 2 | 2 | NA_Cas9_ctrl |
| mock_ctrl4 | NA | 49 | 44 | 4 | 1 | 0 | NA_Cas9_ctrl |
| mock_ctrl5 | NA | 48 | 42 | 3 | 2 | 1 | NA_Cas9_ctrl |
| mock_ctrl6 | NA | 30 | 21 | 4 | 1 | 4 | NA_Cas9_ctrl |
| mock_ctrl7 | NA | 40 | 28 | 2 | 2 | 8 | NA_Cas9_ctrl |
| mock_ctrl1 | NA | 31 | 27 | 1 | 1 | 2 | NA_BE_ctrl |
| mock_ctrl2 | NA | 46 | 34 | 0 | 3 | 9 | NA_BE_ctrl |
| mock_ctrl3 | NA | 28 | 21 | 2 | 0 | 5 | NA_BE_ctrl |

**Table S4 Significance levels of heart rate differences at 21°C, 28°C., and 35°C. P values were assessed with the Wilcoxon test comparing the heart rate of the mutant embryos to the heart rate of the respective mock-injected control embryos.**

| **Gene_group_editor** | **Phenotype** | **p value** | **Temperature** |
| --- | --- | --- | --- |
| adprhl1_Cas9 | normal | 0.049 | 21 °C |
| adprhl1_Cas9 | normal | 0.137 | 28 °C |
| adprhl1_Cas9 | normal | 0.250 | 35 °C |
| adprhl1_evoBE4max | normal | 0.129 | 21 °C |
| adprhl1_evoBE4max | normal | 0.042 | 28 °C |
| adprhl1_evoBE4max | normal | 0.953 | 35 °C |
| blzf1_Cas9 | normal | 0.741 | 21 °C |
| blzf1_Cas9 | normal | 0.984 | 28 °C |
| blzf1_Cas9 | normal | 0.104 | 35 °C |
| blzf1_evoBE4max | normal | 0.009 | 21 °C |
| blzf1_evoBE4max | normal | 0.000 | 28 °C |
| blzf1_evoBE4max | normal | 0.015 | 35 °C |
| btbd1_Cas9 | normal | 0.009 | 21 °C |
| btbd1_Cas9 | normal | 0.033 | 28 °C |
| btbd1_Cas9 | normal | 0.269 | 35 °C |
| btbd1_evoBE4max | normal | 0.002 | 21 °C |
| btbd1_evoBE4max | normal | 0.038 | 28 °C |
| btbd1_evoBE4max | normal | 0.032 | 35 °C |
| gpc5a_Cas9 | normal | 0.053 | 21 °C |
| gpc5a_Cas9 | normal | 0.452 | 28 °C |
| gpc5a_Cas9 | normal | 0.018 | 35 °C |
| gse1_Cas9 | normal | 0.039 | 21 °C |
| gse1_Cas9 | normal | 0.779 | 28 °C |
| gse1_Cas9 | normal | 0.455 | 35 °C |
| irf1_Cas9 | normal | 0.057 | 21 °C |
| irf1_Cas9 | normal | 0.374 | 28 °C |
| irf1_Cas9 | normal | 0.079 | 35 °C |
| pcdh17_Cas9 | normal | 0.710 | 21 °C |
| pcdh17_Cas9 | normal | 0.889 | 28 °C |
| pcdh17_Cas9 | normal | 0.605 | 35 °C |
| phka2_Cas9 | normal | 0.301 | 21 °C |
| phka2_Cas9 | normal | 0.242 | 28 °C |
| phka2_Cas9 | normal | 0.010 | 35 °C |
| phka2_evoBE4max | normal | 0.383 | 21 °C |
| phka2_evoBE4max | normal | 0.465 | 28 °C |
| phka2_evoBE4max | normal | 0.258 | 35 °C |
| ptprd_Cas9 | normal | 0.852 | 21 °C |
| ptprd_Cas9 | normal | 0.844 | 28 °C |
| ptprd_Cas9 | normal | 0.140 | 35 °C |
| ptprd_evoBE4max | normal | 0.607 | 21 °C |
| ptprd_evoBE4max | normal | 0.105 | 28 °C |
| ptprd_evoBE4max | normal | 0.160 | 35 °C |
| rrad_Cas9 | normal | 0.001 | 21 °C |
| rrad_Cas9 | normal | 0.022 | 28 °C |
| rrad_Cas9 | normal | 0.007 | 35 °C |
| rrad_evoBE4max | normal | 0.058 | 21 °C |
| rrad_evoBE4max | normal | 0.113 | 28 °C |
| rrad_evoBE4max | normal | 0.004 | 35 °C |
| sec61a1_Cas9 | normal | 0.195 | 21 °C |
| sec61a1_Cas9 | normal | 0.469 | 28 °C |
| sec61a1_Cas9 | normal | 0.366 | 35 °C |
| zrsr2_Cas9 | normal | 0.008 | 21 °C |
| zrsr2_Cas9 | normal | 0.135 | 28 °C |
| zrsr2_Cas9 | normal | 0.014 | 35 °C |

**Table S5 Knockout target sites.** PAM sites specified in brackets.

| **Name** | **Sequence 5’-3’** |
| --- | --- |
| adprhl1_T1 sgRNA | CTCTGGCGTAGATTTCCAAT[GGG] |
| adprhl1_T2 crRNA | ACAGCCCAAGCTCTCATAAC[AGG] |
| blzf1_T1 sgRNA | TCACGGCTGGGTGATTTGAC[TGG] |
| blzf1_T2 crRNA | GAGTCGAGAGGCCTGTACTG[AGG] |
| btbd1_T1 sgRNA | ACTTATGGGCCGGTATCCGC[TGG] |
| btbd1_T2 crRNA | CAGGCAGGCCCAGCGGATAC[CGG] |
| gpc5a_T1 sgRNA | GATCTGGTTTATCACAGGAT[CGG] |
| gse1_T1 sgRNA | CGAAAGCATAGGGGTTTCCC[AGG] |
| irf1a_T1 sgRNA | CGGGCTTGTCTTTACCCGGA[CGG] |
| pcdh17_T1 sgRNA | ATGCTTAGAGTTACAATAAG[AGG] |
| phka2_T1 sgRNA | GAACGACGTGGTTTGGACAC[TGG] |
| phka2_T2_crRNA | TGTCACCAGGTGAGTCCAGG[AGG] |
| ptprd_T1 sgRNA | ATGAAGACCACAGCCAGCAC [AGG] |
| ptprd_T2 crRNA | CTTTGAACCAAGAAATTTCC[GGG] |
| rrad_T1 sgRNA | TCTGCGGTGTCGTTGCGCGC[AGG] |
| rrad_T2 crRNA | GACACCGCAGACGGGACAAA[CGG] |
| sec61a1_T1 sgRNA | CTACTGGATGAGAGTAATAT[TGG] |
| zrsr2_T1 sgRNA | GAGCTTGTCGATTTGGAGAA[AGG] |

**Table S6 Oligonucleotides for CRISPR sgRNA cloning.**

| **Name** | **Sequence 5’-3’** |
| --- | --- |
| adprhl1_T1_ oligo_F | TAGGCTGGCGTAGATTTCCAAT |
| adprhl1_T1_ oligo_R | AAACATTGGAAATCTACGCCAG |
| blzf1_T1_ oligo_F | TAGGACGGCTGGGTGATTTGAC |
| blzf1_T1_ oligo_R | AAACGTCAAATCACCCAGCCGT |
| btbd1_T1_ oligo_F | TAGGTTATGGGCCGGTATCCGC |
| btbd1_T1_ oligo_R | AAACGCGGATACCGGCCCATAA |
| gpc5a_T1_ oligo_F | TAGGTCTGGTTTATCACAGGAT |
| gpc5a_T1_ oligo_R | AAACATCCTGTGATAAACCAGA |
| gse1_T1_ oligo_F | TAGGAAAGCATAGGGGTTTCCC |
| gse1_T1_ oligo_R | AAACGGGAAACCCCTATGCTTT |
| irf1a_T1_ oligo_F | TAGGGGCTTGTCTTTACCCGGA |
| irf1a_T1_ oligo_R | AAACTCCGGGTAAAGACAAGCC |
| pcdh17_T1_ oligo_F | TAGGGCTTAGAGTTACAATAAG |
| pcdh17_T1_ oligo_R | AAACCTTATTGTAACTCTAAGC |
| phka2_T1_ oligo_F | TAGGACGACGTGGTTTGGACAC |
| phka2_T1_ oligo_R | AAACGTGTCCAAACCACGTCGT |
| ptprd_T1_ oligo_R | AAACGTGCTGGCTGTGGTCTTC |
| ptprd_T1_oligo_F | TAGGGAAGACCACAGCCAGCAC |
| rrad_T1_ oligo_F | TAGGTGCGGTGTCGTTGCGCGC |
| rrad_T1_ oligo_R | AAACGCGCGCAACGACACCGCA |
| sec61a1_T1_ oligo_F | TAGGACTGGATGAGAGTAATAT |
| sec61a1_T1_ oligo_R | AAACATATTACTCTCATCCAGT |
| zrsr2_T1_ oligo_F | TAGGGCTTGTCGATTTGGAGAA |
| zrsr2_T1_ oligo_R | AAACTTCTCCAAATCGACAAGC |

**Table S7 Oligonucleotides used to PCR amplify the target loci.**

| **Name** | **Sequence 5’-3’** |
| --- | --- |
| ptprd_T1_F | CAGACCCACCCACGCTTC |
| ptprd_T1_R | TTCACACCCGCATACTTTGC |
| rrad_T1_F | GCATGGCTCATCTATCCACAGA |
| rrad_T1_R | TGTCTGTGTGGGATGTTTGGT |
| adprhl1_T1_F | ACCCAAACTTGTTGGAAGACAG |
| adprhl1_T1_R | CAGGAGAGCAGTCCACACAC |
| pcdh17_T1_F | GTTACTGGGGAGGTGCGAAC |
| pcdh17_T1_R | GCTCACCAAGGTCAGAGGTC |
| irf1a_T1_F | CATCAACCATTGAGTTGTCACA |
| irf1a_T1_R | TTCTGTGTTTGAGAGCGGGG |
| gpc5a_T1_F | TGCAGAGCTTTTATCAGAAACAGC |
| gpc5a_T1_R | TAAGATGGTTGCTGCAGGGG |
| btbd1_T1_F | AGGACCGATGTACAACTGGC |
| btbd1_T1_R | AGGTCAGTGACTTCCAGCTG |
| gse1_T1_F | AGAGGCTGCCACGTTACAAA |
| gse1_T1_R | GCTGTTGGTGCTCATTCCAG |
| sec61a1_T1_F | GTTTTCTTGAAACTGCAGATCCA |
| sec61a1_T1_R | AAAGCTTGTGAACTACAGATCAAA |
| zrsr2_T1_F | CACTTTGTTTATGAAGTCTGCGT |
| zrsr2_T1_R | GGGAAAAATTGCCTCGAAGTAA |
| blzf1_T1_F | AATGCAACATTCAGGACAGAACA |
| blzf1_T1_R | TGGAGCAGTTGAGTGTAACAGG |
| phka2_T1_F | GTCCAGTTGATGACGGCGAT |
| phka2_T1_R | GAGGACAGAGTGGTGTTTCA |
| adprhl1_T2_ F | GTCCAATCAGCCTAGTTTGGT |
| adprhl1_T2_R | GTTGGGCTACAGGAAGGGAC |
| rrad_T2_F | GTCCCGTTTCCAGTCAACCT |
| rrad_T2_R | TCGTGACCGACATCCTCAAC |
| blzf1_T2_F | CTGTCTGCACTCGAAGCTGA |
| blzf1_T2_R | CCTATTCGGGGTCCTGGAGA |
| btbd1_T2_F | TAAGGGGTCCCGTCTACCTG |
| btbd1_T2_R | TCATGAAGCTGCGTCCAACT |
| phka2_T2_F | TCTGAGGATGCCAGGAGACC |
| phka2_T2_R | AAGATGTTTCAAACACAGTCTGGA |
| ptprd_T2_F | CAGCAGCAGCAAGTTCGAAA |
| ptprd_T2_R | CCTGACAGGTGGCATTGAGA |

**Movie S1**

Swim tunnel assay of HdrR and HO5 individuals. Representative HdrR and HO5 adults in swim tunnel assay at 20m/s water flow. Note the steady swimming behavior at constant speed of the HdrR fish in contrast to the forward pushing and falling back of the HO5 individual during the 10 second movie (played in real time).

**Movie S2**

Embryonic heartbeat of candidate crispants. 10 second movies of embryonic heartbeat at 4 days post fertilization played in real time of unaffected control and *rrad*, *phka2*, *adprhl1*, *ptprd*, *blzf1,* *btbd1* crispants with cardiac phenotypes.

**Movie S3**

Embryonic heartbeat of candidate crispants with cardiac phenotype. 10 second movies of embryonic heartbeat at 4 days post fertilization played in real time of *irf1*, *zrsr2*, *gpc5a*, *pcdh17*, *gse1*, and *sec61a1* crispants with cardiac phenotypes.

**Movie S4**

Embryonic heartbeat of control and *rrad, blzf1 and adprhl1* F0 crispant (top row) and F2 mutants (bottom row). 10 second movies of embryonic heartbeat at 4 days post fertilization played in real time. Note the regular heartbeat in the control embryo versus the different degrees of atrioventricular (AV) block in the F0 crispants and F2 mutants.

**Data S1**

Associated fine mapped regions. Associated fine mapped regions across all temperatures 21°C, 28°C, and 35°C, as well as the variance phenotype (VA); selected candidate genes (green highlighted); see separate Excel file.

**Data S2**

Heart rate data functional validation. Raw data of heartbeat analysis (at 4 dpf) in crispants, editants and mock-injected embryos at 21 °C, 28 °C, and 35 °C; confer separate Excel file.
